# The Metaphase Chromatin Unit: A Novel Unit of Higher-Order Chromosome Organization in Human Mitotic Cells

**DOI:** 10.64898/2026.06.03.729997

**Authors:** Michell Goyal, Ravi Goyal

**Author notes:** Correspondence: Ravi Goyal, MD, PhD.

## Abstract

**Background and Objectives:** The linear relation between human metaphase chromosome length and DNA content has never been rigorously reconciled with modern Hi-C models of mitotic chromatin folding. We tested whether the relation implies a quantitative unit of mitotic chromosome organization.

**Methods:** We pooled metaphase lengths for 24 human chromosomes from five cytogenetic studies of cultured peripheral lymphocytes and regressed length against base-pair content from GRCh38 and T2T-CHM13 v2.0. Pre-specified analyses comprised ordinary least-squares and power-law fits, arm-level decomposition, and reconciliation with the Gibcus 2018 helical loop-array model. Orthogonal validation used Rao 2014 Hi-C boundary counts and Pope 2014 Repli-Seq.

**Results:** Length scales linearly with DNA content as L (μm) = 0.0329 x Mb + 0.043 (R-squared = 0.998, power-law exponent 0.98 +/- 0.01), with a cross-karyotype compaction density of 33.4 +/- 1.0 nm/Mb. T2T-CHM13 reanalysis identifies satellite over-condensation; the arm-level residual correlates with p-arm fraction (r = 0.65, p = 5.6e-4). We propose an operational Metaphase Chromatin Unit (MCU) as a quantitative scaling unit (not a discrete structural quantum, as quantization tests are negative): 1 MCU = 7.6 Mb DNA = 0.25 μm axial length, numerically corresponding to one Gibcus late-metaphase helical turn; the human haploid genome scales to 406 MCUs. The pairwise megabase-shift slope of 32.75 nm/Mb (R-squared = 0.996) and the approximately 6 Mb Abbe optical detection threshold correctly classify 9 of 9 microdeletion syndromes as karyotype-visible versus FISH-required.

**Conclusions:** The MCU provides a unified quantitative unit for human mitotic chromosome organization, integrating cytogenetic, Hi-C, polymer-biophysical, and clinical scales.

## 1. Introduction

The mammalian genome is hierarchically packaged across several orders of length scale, from the 2 nm DNA duplex to metaphase chromatids of variable dimensions (approximately 500 to 700 nm wide and 1.6 to 8.2 μm long for the 24 human chromosomes, with values dependent on cell-cycle timing and preparation conditions) [1, 2]. Interphase Hi-C established that mammalian chromatin partitions into topologically associating domains (TADs) with a median size of approximately 880 kb [3] and into smaller contact domains and loops of median 185 kb with approximately 10,000 loops genome-wide [4]. TAD boundaries are demarcated by CTCF and cohesin [5]. During mitosis the interphase pattern is replaced by a uniform, helically arranged loop array generated by condensin-driven loop extrusion [6–8]. Gibcus and colleagues showed that the late-metaphase helical turn contains approximately 12 Mb of DNA in ∼150 loops at an axial spacing of 200 to 250 nm [7].

Alternative and complementary structural models of the mitotic chromatid include the classical unit-fiber and higher-order coiling models [9], the recently reviewed conservation of helical chromonema coiling across eukaryotes [10], and the multilayer (stacked-plate) chromatin model developed from atomic force microscopy and cryo-electron microscopy studies of chromatin emanated from chromosomes [11–13], in which chromatids are stacks of approximately 6 nm thick planar chromatin layers each containing approximately 0.5 Mb of DNA and packed to a volumetric density of approximately 0.17 Gb per cubic micrometre, with cytogenetic bands, sister chromatid exchanges, and cancer translocations oriented orthogonal to the chromatid axis [12]. Across species, chromosome length has been shown to scale with DNA content approximately as the square root of Mb across more than 600 vertebrates and angiosperms [14]; within a single species where chromatid diameter is approximately constant, length is therefore expected to scale nearly proportionally with DNA content.

We emphasize that metaphase is not a single instantaneous state but a range of compaction states spanning prometaphase to early anaphase; karyometric measurements from colchicine-arrested lymphocyte preparations pool observations across a spread of individual cell-cycle time-points, and karyotypic heterogeneity between and within cells is well documented [15]. All axial-length values reported in this work should therefore be understood as population averages over that spread, not as instantaneous single-cell values. Furthermore, the 33 nm/Mb constant is a chromosome-scale population average that does not describe local intra-chromosomal DNA packing, which varies markedly between GTG-dark and GTG-light bands as documented by Claussen and colleagues, and is naturally captured at the single-band scale by the multilayer chromatin model [16].

A first-principles consequence of any such model is that the linear length of the mitotic chromatid should be a smooth, near-proportional function of DNA content with a per-Mb axial density predictable from the helical loop geometry. Classical cytogenetic measurements of relative chromosome length hinted at this proportionality but predated finished reference assemblies and were never formally regressed against base-pair counts [17–19].

### Glossary

Throughout this work we use three distinct quantitative concepts. The Metaphase Chromatin Unit (MCU) is an operational scaling unit, not a discrete structural quantum: 1 MCU is defined as 7.6 Mb of DNA whose compaction at 33.4 nm/Mb yields 0.25 μm of metaphase axial length, numerically equal to one Gibcus late-metaphase helical turn. The cytogenetic data are consistent with smoothly proportional compaction without discrete quantization (Section 3.2); the MCU therefore functions as a quantitative benchmark for translation between Mb, axial length, and Hi-C scales, not as a separable physical feature of the mitotic chromosome. The megabase-shift relation is the related pairwise heuristic that adding or removing approximately 6 to 8 Mb of DNA shifts a chromosome by approximately one MCU of metaphase length. The approximately 6 Mb cytogenetic detection threshold is the Abbe diffraction limit of light microscopy translated through the 33 nm/Mb scaling; it is numerically similar to but conceptually distinct from the 7.6 Mb MCU.

In this report we pool mean metaphase lengths for 24 human chromosomes from five independent cytogenetic studies, regress length against finished base-pair content from GRCh38 and T2T-CHM13 v2.0, decompose the residual structure into arm-level and chromatin-state components, reconcile the result quantitatively with the Hi-C helical loop-array model, and define the MCU as an operational quantitative scaling unit at the helical-turn scale. Cross-species predictions, chromothripsis applications, synthetic biology design rules, microdeletion syndrome classification, polymer physics, and an interactive supplementary calculator are provided in the Supplementary Material.

## 2. Materials and Methods

### 2.1. Pre-specified primary analyses

We pre-specified five primary analyses before unblinding: (i) ordinary least-squares regression of metaphase length on Mb under GRCh38; (ii) the same regression under T2T-CHM13 v2.0; (iii) arm-level decomposition of the residual using cytoband centromere positions; (iv) the helical-turn reconciliation under the Gibcus 2018 axial spacing; (v) the pairwise (276-pair) inter-chromosomal regression. All other analyses, including the chromHMM cross-validation, Repli-Seq comparison, cross-species predictions, microdeletion syndrome classification, polymer-physics framing, and chromothripsis applications, are reported as exploratory and are detailed in the Supplementary Material.

### 2.2. Cytogenetic length data

Mean lengths of human metaphase chromosomes 1 to 22, X, and Y were taken from five independent published studies of cultured peripheral lymphocytes [6, 19–22]. We used the arithmetic mean across studies as the point estimate and the inter-study sample standard deviation as a measurement-noise estimate. The five studies used standard light microscopy of colchicine-arrested peripheral lymphocyte preparations with an inherent optical resolution of approximately 200 nm (Abbe diffraction limit at typical NA 1.3 to 1.4 and visible wavelengths); the ∼50 nm inter-study SD per chromosome reflects averaging over hundreds of individual cells per study, and is at or slightly below the single-image optical resolution of these preparations. Lengths ranged from 1.576 μm (chromosome 21) to 8.208 μm (chromosome 1).

### 2.3. Reference DNA content

Base-pair counts were taken from (i) the GRCh38 primary assembly [23], and (ii) the T2T-CHM13 v2.0 assembly (GenBank GCA_009914755.4) [24]. Centromere positions for the arm-level analysis were obtained from the UCSC cytoBand track for hg38 (acen entries).

### 2.4. Statistical analysis

We performed OLS linear regression of mean length on Mb, computed slope, intercept, R-squared, slope 95 percent confidence interval, a zero-intercept fit, a log-log power-law fit, and per-chromosome residuals. Residual quantization at 0.23 μm was tested by computing the residual standard deviation from the nearest 0.23 μm multiple. The arm-level analysis decomposed each chromosome into p-arm Mb and q-arm Mb using cytoband centromere positions and tested the Pearson correlation between p-arm fraction and the GRCh38 length residual. Pairwise inter-chromosomal differences (276 chromosome pairs) were regressed delta-L on delta-Mb. Analyses used Python 3.10 with NumPy 1.26 and SciPy 1.13. Code and data are deposited (see Data Availability).

## 3. Results

### 3.1. Length scales linearly with DNA content at 33.4 nm/Mb

Across all 24 chromosomes, metaphase length increases linearly with DNA content (Figure 1A). The OLS fit is L (μm) = 0.03291 (+/- 0.00031) x Mb + 0.043 (+/- 0.044), R-squared = 0.998, p = 2.3e-31, 95 percent CI for the slope [0.0323, 0.0335] μm/Mb. Forcing the regression through the origin gives a slope of 0.03319 μm/Mb with R-squared = 0.998, indicating the intercept is not statistically distinguishable from zero. A power-law fit yields an exponent of 0.981 +/- 0.012 (R-squared = 0.997), within 2 standard errors of unity. Per-chromosome residuals from the linear fit are shown in Figure 1B; per-chromosome compaction density is 33.4 +/- 1.0 nm/Mb with a coefficient of variation of 2.9 percent (Figure 1C). The X chromosome (32.95 nm/Mb) and Y chromosome (32.26 nm/Mb) sit within one SD of the autosomal mean. Acrocentric chromosomes 13, 14, and 15 have somewhat lower compaction densities (31.30, 31.84, 32.28 nm/Mb), reflecting shorter measured lengths relative to total DNA content; chromosome 19, the most gene-dense autosome, has the highest at 36.10 nm/Mb.

**Figure 1.**
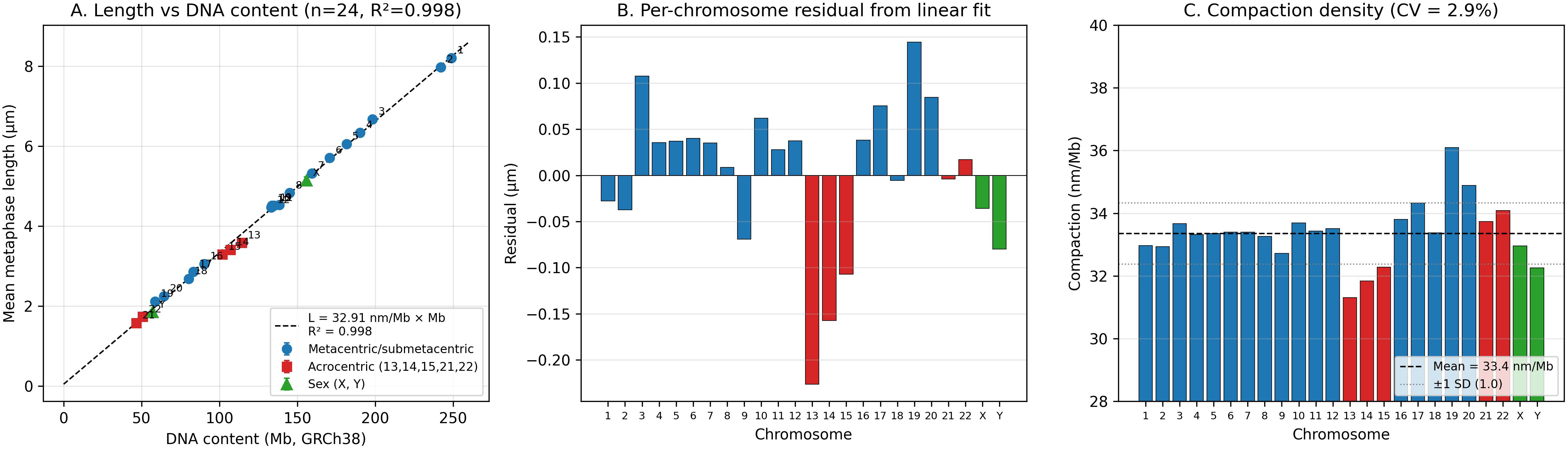
(A) Mean metaphase length vs DNA content for 24 human chromosomes (GRCh38). Circles = metacentric/submetacentric; squares = acrocentric (13, 14, 15, 21, 22); triangles = sex chromosomes. Dashed line: OLS fit, R-squared = 0.998. (B) Per-chromosome residuals from the linear fit. (C) Compaction density (nm of metaphase per Mb of DNA) across chromosomes, mean 33.4 nm/Mb, CV 2.9 percent.

### 3.2. No statistical support for a discrete megabase folding unit

A previously suggested interpretation of the linear scaling is that the chromosome is built from a discrete number of folding domains of approximately 6.8 Mb each, contributing approximately 0.23 μm per domain [6]. The chr 1 versus chr 2 length difference of 0.232 μm for 6.76 Mb of DNA appears to support this. Across all 24 chromosomes, however, the residual standard deviation from the nearest 0.23 μm multiple (0.071 μm) is essentially identical to the residual SD from the smooth linear fit (0.084 μm); the same holds for 0.25 μm and 0.32 μm quantizations. No specific quantization length improves the fit over continuous proportionality. The data are consistent with a smoothly proportional compaction at the whole-chromosome scale. We note the resolution limit of this test: at the observed 33 nm/Mb, a hypothetical 500 kb folding unit would correspond to approximately 17 nm of axial length, well below both the inter-study SD (∼50 to 90 nm) and the ∼200 nm optical resolution of the light-microscopy preparations. Sub-megabase quantization, including the approximately 0.5 Mb per 6 nm chromatin layer proposed by the Daban multilayer model [11–13], is therefore not testable by karyometric measurement alone and would require higher-resolution structural methods.

### 3.3. T2T-CHM13 reanalysis identifies satellite over-condensation

Repeating the regression using T2T-CHM13 v2.0 Mb values yields a slope of 0.03286 μm/Mb and R-squared = 0.996, only marginally different from GRCh38. The residual structure changes substantially (Figure 2A, Supplementary Table S1). Chromosomes that received the largest T2T heterochromatin additions show large new negative residuals: chromosome 9 (gained 12.2 Mb of pericentromeric satellite) acquires a residual of -0.43 μm versus -0.07 μm on GRCh38; chromosome Y (gained 5.2 Mb) reaches -0.22 μm versus -0.08 μm; chromosome 16 (gained 6 Mb of satellite) reaches - 0.12 μm versus +0.04 μm. The 12.2 Mb chr 9 satellite block contributes negligible incremental length beyond the inter-study noise floor of approximately 53 nm, bounding the satellite compaction density at <= 4.3 nm/Mb, that is, at least 8-fold more compact than the chromosome-wide mean of 33.4 nm/Mb. T2T-CHM13 therefore provides a cytogenetic-scale inference consistent with marked satellite over-condensation.

**Figure 2.**
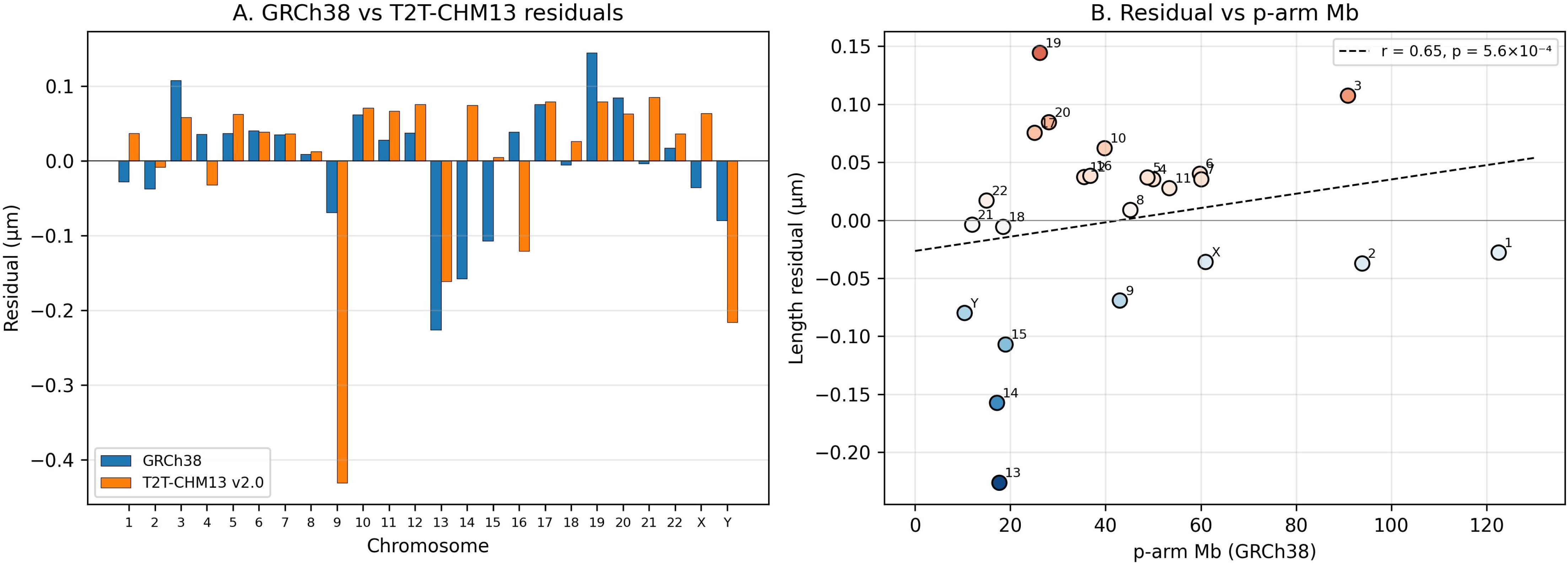
(A) Per-chromosome residual using GRCh38 (blue) vs T2T-CHM13 v2.0 (orange). Chromosomes 9, 16, and Y, which received the largest T2T heterochromatin additions, show large new negative residuals. (B) Length residual vs p-arm Mb; Pearson r = 0.65, p = 5.6e-4. Acrocentric chromosomes 13 to 15 cluster at low p-arm Mb with negative residuals.

### 3.4. Arm-level decomposition: p-arm fraction predicts the residual

Decomposing each chromosome into p-arm Mb and q-arm Mb using GRCh38 cytoband centromere positions, the length residual correlates strongly with p-arm fraction (Pearson r = 0.65, p = 5.6e-4; Figure 2B). However, this arm-level association is attenuated under the T2T-CHM13 v2.0 reanalysis (Supplementary Section S2), indicating that the GRCh38 p-arm signal should be interpreted cautiously and may partly reflect incomplete acrocentric short-arm representation in GRCh38. The five chromosomes with the smallest p-arm fraction (13, 14, 15, Y, 18) include three of the four largest negative residuals; chromosomes with intermediate-to-large p-arm fraction (16, 19, 20, 17) include the largest positive residuals. This is consistent with satellite over-condensation of the acrocentric short arms (Section 3.3). The chromHMM-based analysis (active chromatin r = +0.48; heterochromatin r = -0.50; Supplementary Section S3) and Pope 2014 Repli-Seq comparison (Supplementary Section S4) corroborate this interpretation. We emphasize that the 33 nm/Mb constant is a chromosome-scale average and not a locally uniform statement about DNA packing along a single chromosome. Well-documented differences in local compaction between GTG-dark bands (denser) and GTG-light bands (more decondensed) at the single-chromosome level have been reported by Claussen and colleagues, and are compatible with the multilayer chromatin model of Daban and colleagues, in which each of stacked approximately 6 nm planar layers can contain different local DNA densities and in which chromosome bands, sister chromatid exchanges, and translocations are systematically oriented orthogonal to the chromatid axis as expected for stacked planar layers [11–13, 16]. Consistent with these observations, fluorescence in situ hybridization with probes at equal genomic spacing along a single chromosome typically shows unequal axial spacing depending on local band and chromatin-state context. The MCU framework therefore describes population-averaged chromosome-scale length as a linear function of DNA content, not local intra-chromosomal DNA density.

### 3.5. Quantitative reconciliation with the Hi-C helical loop-array model

The Gibcus 2018 helical loop-array model predicts a per-turn axial density (Figure 3A) of approximately 67 nm/Mb in early metaphase (3 Mb per 0.20 μm turn) and approximately 21 nm/Mb in late metaphase (12 Mb per 0.25 μm turn) [7]. Our cross-chromosome measurement of 33.4 nm/Mb sits between these and is closest to a mid-prometaphase regime, consistent with the timing of colchicine-arrested lymphocyte preparations that underlie the cytogenetic data. The 200 to 250 nm axial spacing of helical turns reported by Gibcus et al. (2018) in synchronised DT40 chicken cells is preserved as a length scale in our data; the DNA content per unit axial length, however, is lower in our karyometric measurements (7.6 Mb per 250 nm) than in the late-metaphase DT40 measurements of Gibcus et al. (12 Mb per 250 nm turn). Under the helical loop-array framework, this discrepancy is consistent with a less-condensed mid-prometaphase state characteristic of colchicine-arrested lymphocyte preparations, or with cell-type differences in condensin loading between chicken DT40 and human lymphocytes. The MCU is therefore not equivalent to a Gibcus late-metaphase helical turn; it corresponds to the same axial length scale but at a different DNA content per unit length, which is the physically expected observation for a less-condensed metaphase state. The 33 nm/Mb axial density corresponds to approximately 6.1 nm per Rao 2014 contact domain, approximately 29 nm per Dixon 2012 TAD, and approximately 13 nm per Gibcus 2018 prometaphase outer loop; full real per-chromosome Hi-C boundary regression against length yields R-squared = 0.84 (Figure 3B; Supplementary Section S5).

**Figure 3.**
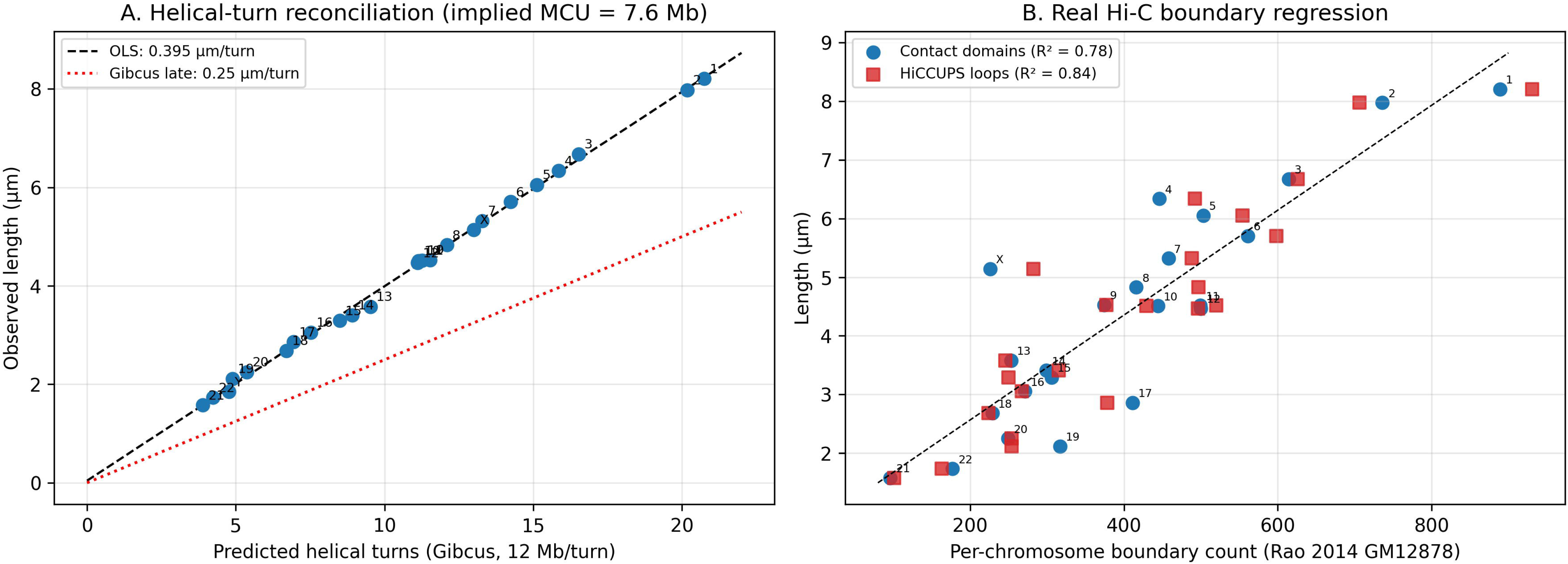
(A) Observed metaphase length plotted against predicted helical-turn count under the Gibcus late-metaphase parameter (12 Mb per turn). Dashed line: OLS fit (0.395 μm per turn). Dotted line: Gibcus 0.25 μm per turn. The implied DNA per turn under the human data is 7.6 Mb (= 1 MCU). (B) Real per-chromosome Hi-C boundary regression using Rao 2014 GM12878 HiCCUPS loops (red) and Arrowhead contact domains (blue). Slope 8.8 and 8.9 nm per boundary; R-squared 0.84 and 0.78.

### 3.6. The megabase-shift relation across all chromosome pairs

Computing the 276 pairwise inter-chromosomal differences in length and Mb (Figure 4A; the corresponding quantization test for Section 3.2 is shown in Figure 4B), the pairwise slope is 32.75 nm/Mb (95 percent CI [32.49, 33.00]; R-squared = 0.996). Pairwise differences are shown symmetrically about zero because every chromosome pair (i, j) yields both a positive difference (i > j) and a negative difference (j > i); the sign carries no biological meaning and reflects only the arbitrary ordering of the two chromosomes in each pair. The pairwise analysis is complementary to the direct axial-length regression: it tests whether the compaction density is constant across chromosomes of different absolute sizes, and therefore whether the linear scaling holds when the largest and smallest chromosomes are directly compared rather than jointly fitted. The original chr 1 versus chr 2 observation (6.76 Mb, 0.232 μm) lies on this line. The pairwise relation generalizes the chr 1-chr 2 observation to a generic statement applicable between any two chromosomes: adding or removing approximately 6 to 8 Mb of DNA shifts metaphase length by approximately 0.20 to 0.27 μm, that is, approximately one MCU.

**Figure 4.**
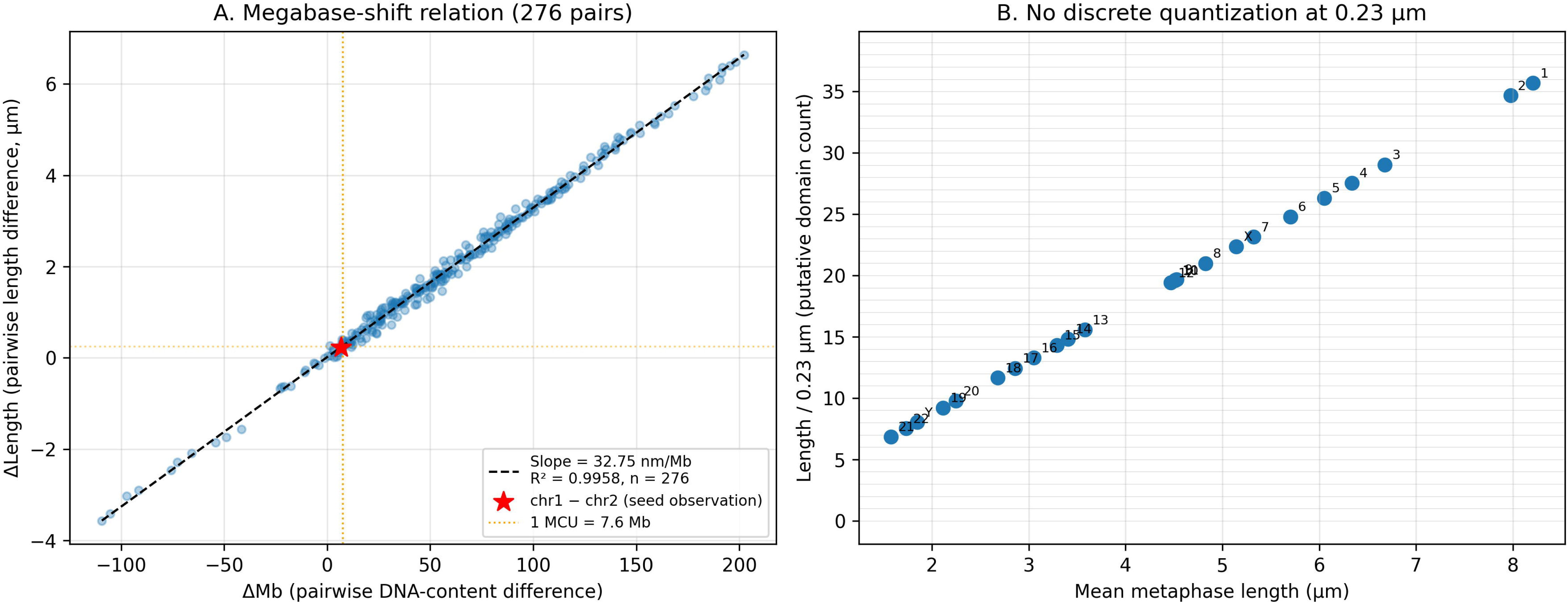
(A) Pairwise inter-chromosomal scaling: each point is one of 276 chromosome pairs. Slope 32.75 nm/Mb, R-squared = 0.996. Red star: the original chr 1 vs chr 2 observation (6.76 Mb, 0.232 μm). Orange dotted lines: 1 MCU at 7.6 Mb / 0.25 μm. (B) Quantization test: putative domain count (length divided by 0.23 μm) does not cluster at integer values, indicating no detectable quantization at the 0.23 μm scale.

### 3.7. The Metaphase Chromatin Unit (MCU)

Building on the helical-turn reconciliation of Section 3.5, we propose the MCU as an operational quantitative unit of human mitotic chromosome organization. Importantly, the MCU is not a discretely separable structural object in the chromosome (Section 3.2 finds no statistical support for quantization at any specific length scale, including 7.6 Mb); rather, it is a quantitative benchmark obtained by treating the empirical 33.4 nm/Mb linear compaction as if it were partitioned into integer blocks whose size matches the published Gibcus late-metaphase helical-turn dimensions. Under this operational definition: 1 MCU = 7.6 Mb of DNA = 0.25 μm of metaphase axial length, numerically corresponding to one Gibcus late-metaphase helical turn = approximately 41 Rao 2014 contact domains = approximately 9 Dixon 2012 TADs = approximately 150 Gibcus prometaphase mitotic loops. The human haploid genome contains 3,088 Mb / 7.6 Mb = 406 MCUs and produces a total mitotic axial length of 406 x 0.25 = 101.6 μm, in close agreement with the pooled observed total of 102.7 μm. Individual chromosomes range from 6.1 MCUs (chromosome 21) to 32.8 MCUs (chromosome 1). Because the MCU is defined as an operational scaling unit rather than a discrete molecular architecture, it is not an additional structural level of the chromatin hierarchy; it is a chromosome-scale reference distance that happens to be numerically similar to the axial spacing of published mitotic helical turns and to the height of approximately 15 stacked chromatin layers under the Daban multilayer model [12]. Notably, the high in vivo DNA concentration of metaphase chromatids (approximately 0.17 Gb per cubic micrometre) that underlies our observed 33 nm/Mb axial constant is more naturally explained by the multilayer model, in which nucleosome-nucleosome interactions between adjacent planar layers pack DNA densely, than by extended-fibre or loop-extrusion-only models [11, 13]; a physically consistent reconciliation, suggested by Daban and colleagues, is that Hi-C-detected nested loops are compacted into the chromatin layers seen in microscopy. Figure 5 therefore places the MCU as an operational reference annotation alongside the structural levels rather than at Level 9 of the hierarchy itself. The MCU sits alongside the compaction hierarchy of B-form DNA (Level 1), nucleosome (L2), 10 nm nucleosomal fiber (L3), the disputed 30 nm chromatin fiber (L4, whose existence in vivo remains debated; retained here as a historically named level rather than a confirmed architecture), Rao contact domain (L5), prophase and prometaphase loops (L6, L7), Dixon TAD (L8), to MCU (L9) and whole chromatid (L10) (Figure 5, Supplementary Table S3). Cumulative compaction at the MCU level is approximately 10,000-fold linear.

**Figure 5.**
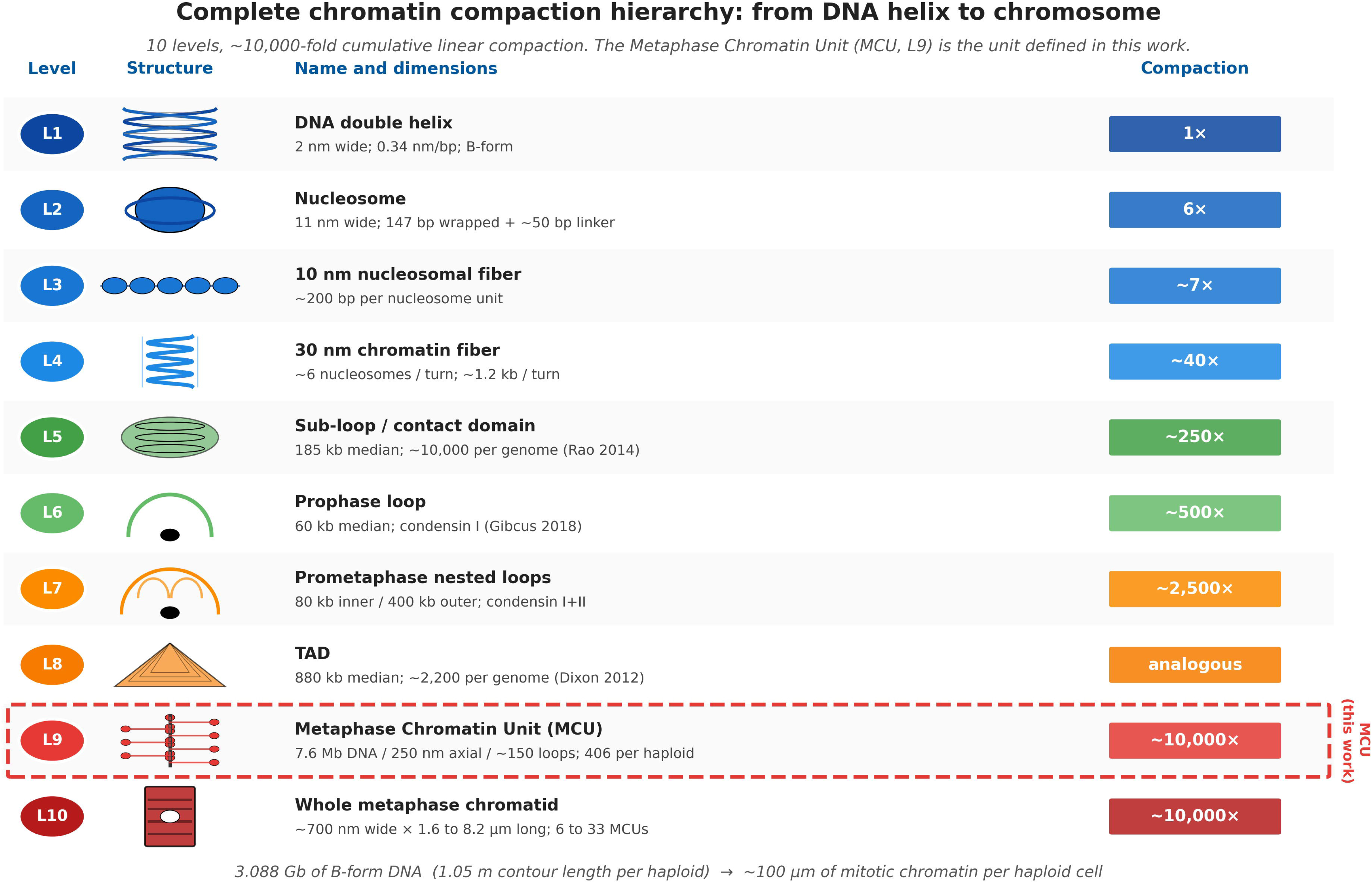
Complete chromatin compaction hierarchy from B-form DNA (L1) to whole metaphase chromatid (L10), with structural cartoons for each level. The MCU (L9, red-bordered) closes the previously missing quantitative gap between mitotic loops (L6 to L7) and whole chromosomes.

### 3.8. Clinical CNV detectability and the approximately 6 Mb optical-resolution threshold

A practical consequence of the 33.4 nm/Mb constant is that the minimum cytogenetically detectable copy number variant can be predicted from optical diffraction theory. The Abbe diffraction limit at NA 1.4 and wavelength 550 nm is 196 nm. At 33.4 nm/Mb, this corresponds to a minimum visible deletion or duplication of 5.9 Mb (Rayleigh equivalent 7.2 Mb), which we summarize as the approximately 6 Mb detection threshold (numerically similar to but conceptually distinct from the 7.6 Mb MCU defined in Section 3.7: the 6 Mb threshold is the optical diffraction limit, whereas the MCU is the helical-turn unit). Standard 400-band karyotype empirically resolves deletions of 5 to 10 Mb and high-resolution 850-band prometaphase karyotype resolves 3 to 5 Mb [25]. The framework correctly classifies 9 of 9 well-characterized contiguous-gene microdeletion syndromes as karyotype-visible vs FISH-required using this single 6 Mb threshold: Williams (1.6 Mb), 22q11.2 typical DiGeorge (3 Mb), 1p36 (3.5 Mb), Smith-Magenis (3.7 Mb), Angelman/PWS (5.6 Mb) are FISH-required; 22q11.2 atypical (6.5 Mb), Jacobsen (11 Mb), Wolf-Hirschhorn (25 Mb), Cri-du-chat (30 Mb) are karyotype-visible (full table in Supplementary Section S6). Direct empirical testing of integer-multiple clustering of recurrent microdeletion sizes at chromatin domain scales (185 kb, 880 kb, MCU multiples) returned a negative result; recurrent CNV sizes are positioned by segmental duplications, not by chromatin domain boundaries (Supplementary Section S7). The 33 nm/Mb framework therefore predicts the cytogenetic detectability of CNVs of any size, without predicting their absolute size distribution. Figure 6 collates the full MCU framework into a single-page dictionary translating between Mb, metaphase length, Hi-C unit counts, polymer-biophysical parameters, evolutionary tests, and clinical CNV detectability for everyday reference use.

**Figure 6.**
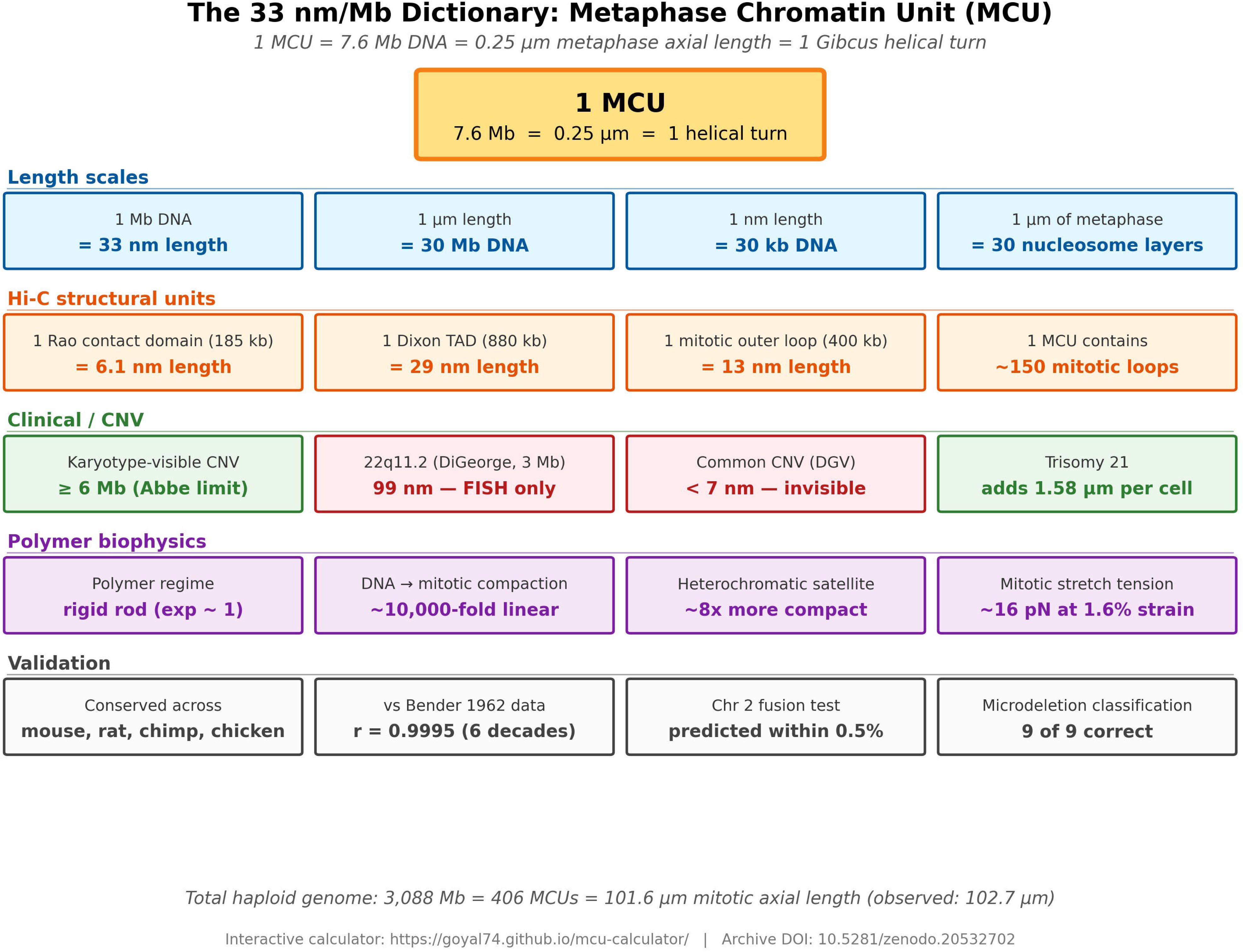
The 33 nm/Mb dictionary: integrated single-page reference for the MCU framework, translating between cytogenetic, Hi-C, polymer-biophysical, clinical, and evolutionary scales. An interactive web-based MCU calculator (Supplementary File MCU_Calculator.html) is live at https://goyal74.github.io/mcu-calculator/ and archived at https://doi.org/10.5281/zenodo.20532702.

## 4. Discussion

Human metaphase chromosome length is a near-perfect linear function of DNA content (R-squared = 0.998) with a cross-karyotype compaction density of 33.4 +/- 1.0 nm/Mb that varies by less than 3 percent across the karyotype. Caspersson and colleagues already noted in the 1970s that relative chromosome lengths track relative DNA content [18], but the precision of the fit with finished assembly base-pair counts and pooled length data from five studies, and the explicit reframing as a quantitative constraint on Hi-C structural models, are new. Four main implications follow.

### 4.1. Reconciliation with the helical loop-array model

The Gibcus 2018 pathway model predicts per-turn axial densities of approximately 67 nm/Mb (early metaphase) and approximately 21 nm/Mb (late metaphase). Our cross-chromosome measurement of 33 nm/Mb sits between these, consistent with mid-prometaphase colchicine-arrested lymphocyte preparations. The 250 nm axial spacing of Gibcus 2018 helical turns is preserved as a length scale in our data, but at 7.6 Mb per 250 nm rather than the 12 Mb per 250 nm reported for late-metaphase DT40 chicken cells. This is consistent with our data being from colchicine-arrested lymphocytes in a less-condensed prometaphase state; the MCU is defined operationally at the 7.6 Mb / 250 nm axial length observed in human lymphocyte preparations. A physically consistent reconciliation of the helical loop-array and multilayer frameworks, suggested by Daban and colleagues, is that the long nested loops detected in Hi-C experiments could be compacted into the planar chromatin layers observed in microscopy studies; under this view the two models are not competing alternatives but complementary descriptions at different structural scales, with nucleosome-nucleosome interactions between adjacent layers providing the physical basis for the observed high in vivo DNA concentration [12, 13]. Our chromosome-scale linear scaling at 33.4 nm/Mb is consistent with either framework at the macroscopic level; distinguishing them requires nanoscale structural methods rather than karyometry.

### 4.2. The MCU as a unifying operational unit

The MCU integrates DNA content, axial length, helical-turn structure, Hi-C boundary counts, and clinical detection thresholds into a single integer arithmetic. We emphasize that the MCU is an operational scaling unit, not a discrete structural quantum, since our quantization tests are negative (Section 3.2): the cytogenetic data are equally well described by a smooth proportional compaction. The MCU is therefore a useful translation benchmark, analogous to how the helical pitch of B-form DNA (3.4 nm per 10 bp) is a convenient unit even though the helical structure is continuous, not stepwise. The compaction hierarchy L1 (DNA duplex) through L10 (whole chromatid) is a structural hierarchy of chromatin architectures; the MCU is not itself one of these architectures and does not close the hierarchy. Instead, it provides a quantitative reference distance annotated alongside the hierarchy, in the same way that the pitch of B-form DNA (3.4 nm per 10 bp) is annotated alongside the L1 duplex without being a separate level. Under the alternative multilayer model of Daban, the same 7.6 Mb / 250 nm length scale corresponds to approximately 15 stacked planar chromatin layers. In MCU units, the chr 1 versus chr 2 difference is one MCU, trisomy 21 adds 6.1 MCUs of extra metaphase chromatin, and the cytogenetic CNV detection threshold is approximately 1 MCU.

### 4.3. The megabase shift and clinical resolution

The pairwise megabase-shift relation (slope 32.75 nm/Mb, R-squared = 0.996 across 276 pairs) provides a simple heuristic: every ∼7 Mb adds or removes ∼0.25 μm of metaphase length. Combined with the Abbe diffraction limit, this predicts the historically observed 6 to 10 Mb cytogenetic detection threshold and correctly classifies 9 of 9 microdeletion syndromes. Recurrent CNV sizes are not quantized at chromatin domain multiples (Supplementary Section S7); they are positioned by segmental duplications. The MCU therefore predicts detectability without predicting CNV size distribution.

### 4.4. Limitations

The five pooled cytogenetic studies span 1964 to 2013 with varying fixation protocols; the cross-study CV is 1.6 percent but inter-laboratory bias cannot be excluded. The arm-level analysis uses GRCh38 centromere positions; T2T-CHM13 acrocentric p-arms are smaller and reduce the arm-level correlation (Supplementary Section S2). Cross-species predictions (Supplementary Section S8) apply the human slope to non-human Mb values and are framed as falsifiability checks rather than independent slope tests. Direct future tests of the MCU would benefit from matched Hi-C and cytometric measurements in the same primary preparations.

### 4.5. Relation to cross-species scaling laws

Kramer and colleagues have compiled metaphase chromosome dimensions across more than 600 species of vertebrates and angiosperms and reported that across species chromosome length scales approximately as Mb to the 1/2 power, width as Mb to the 1/4 power, and volume as Mb to the 1 power [14]. Our intra-human exponent of 0.98 does not contradict this cross-species finding: within a single species chromatid diameter varies little (D approximately 0.4 to 0.6 μm across the 24 human chromosomes), so length scales approximately as Mb^1; across species, D scales as Mb^(1/4) and length as Mb^(1/2), which recovers the observed volume L Mb^1 relationship. The MCU framework is therefore complementary to the Kramer scaling laws: those laws describe how chromosome shape scales across species; we quantify the per-Mb axial constant within a single species and use it as an operational bridge to Hi-C, polymer-biophysical, and clinical CNV scales.

## 5. Conclusions

Human metaphase chromosome length is a near-perfect linear function of DNA content with a cross-karyotype compaction density of 33.4 nm/Mb. We propose an operational Metaphase Chromatin Unit (MCU) = 7.6 Mb of DNA = 0.25 μm of metaphase axial length. This axial length is numerically equal to the pitch of Gibcus 2018 helical turns, though at a lower DNA content per unit pitch than the fully condensed late-metaphase state (12 Mb per turn) reported in DT40 chicken cells, consistent with mid-prometaphase colchicine-arrested lymphocyte preparations; the MCU is also numerically equal to the height of approximately 15 stacked chromatin layers under the Daban multilayer model. In Hi-C terms, one MCU corresponds to approximately 41 Rao 2014 contact domains and approximately 150 prometaphase mitotic loops, with the human haploid genome containing 406 MCUs. The MCU is an operational reference scale annotated alongside (not part of) the structural chromatin hierarchy, and provides a single integer arithmetic translation between cytogenetic axial length, Hi-C boundary counts, polymer-biophysical parameters, and clinical CNV detection thresholds. The MCU is a chromosome-scale population average; it does not describe local intra-chromosomal DNA packing, which varies between GTG-dark and GTG-light bands and is more accurately represented at the single-band scale by the multilayer chromatin model of Daban and colleagues [12], in which mitotic chromatids consist of stacked 6 nm planar chromatin layers packed by nucleosome-nucleosome interactions. Under a reconciliation proposed by Daban, the long nested loops observed in Hi-C studies are compacted into these chromatin layers, so the loop-array and multilayer frameworks are complementary descriptions of the same physical chromatid at different structural scales rather than competing alternatives. The framework is consistent with, and complementary to, the cross-species scaling laws of Kramer and colleagues. The approximately 6 Mb optical detection threshold derived from the Abbe limit, which is numerically close to one MCU but conceptually distinct, correctly classifies 9 of 9 microdeletion syndromes as karyotype-visible versus FISH-required. An interactive MCU calculator is provided as Supplementary Material.

## Supporting information

Online Calculator

## Supplementary Materials

The following supplementary materials are available online. Supplementary Document: extended analyses in 10 sections, four tables, and four figures. Supplementary Table S1: pooled per-chromosome data for 24 human chromosomes with GRCh38 Mb, T2T-CHM13 v2.0 Mb, inter-study standard deviation, compaction density, and residuals from both assemblies. Supplementary Table S2: clinical microdeletion syndrome classification (9 syndromes; predicted versus empirical karyotype detection). Supplementary Table S3: complete chromatin compaction hierarchy (10 levels from B-form DNA to whole chromatid). Supplementary Table S4: aneuploidy length predictions. Supplementary Figure S1: multi-species cross-validation (mouse, rat, chimp, chicken). Supplementary Figure S2: clinical and polymer-physics applications (microdeletion syndromes, aneuploidy, chromothripsis, polymer scaling). Supplementary Figure S3: orthogonal molecular validation (Pope 2014 Repli-Seq, Roadmap E116 chromHMM, variance partitioning, Bender 1962). Supplementary Figure S4: test for integer-multiple clustering of recurrent microdeletion sizes (negative result). Supplementary File MCU_Calculator.html: interactive web-based unit calculator (live at https://goyal74.github.io/mcu-calculator/; archived at https://doi.org/10.5281/zenodo.20532702). Source data: Manuscript_Chromosome_Domain.xlsx.

## Author Contributions

Conceptualization, R.G.; data curation, M.G. and R.G.; formal analysis, M.G. and R.G.; methodology, R.G.; visualization, M.G. and R.G.; writing-original draft, M.G. and R.G.; writing-review and editing, R.G. All authors have read and agreed to the published version of the manuscript.

## Funding

This research received no external funding.

## Institutional Review Board Statement

Not applicable. This study used only previously published, de-identified cytogenetic data and public reference assemblies.

## Informed Consent Statement

Not applicable.

## Data Availability Statement

All primary data are presented in the manuscript and in Supplementary Table S1 (Manuscript_Chromosome_Domain.xlsx). Reference assemblies: GRCh38 (NCBI), T2T-CHM13 v2.0 (NCBI GCA_009914755.4), GRCm39, mRatBN7.2, panTro6, galGal6. Public datasets used: Rao 2014 (GEO GSE63525), Pope 2014 Repli-Seq (GEO GSE53984), Roadmap Epigenomics E116 chromHMM 15-state segmentation. Analysis code and the interactive MCU calculator are publicly available at https://github.com/goyal74/mcu-calculator and live at https://goyal74.github.io/mcu-calculator/. The calculator is permanently archived on Zenodo (DOI: 10.5281/zenodo.20532702; https://doi.org/10.5281/zenodo.20532702). The MCU calculator is also provided as Supplementary File MCU_Calculator.html.

## Conflicts of Interest

The authors declare no conflict of interest.

## Notes

https://github.com/goyal74/mcu-calculator

https://doi.org/10.5281/zenodo.20532702

### Competing Interest Statement

The authors have declared no competing interest.

### Summary of Updates

Manuscript substantially revised following community feedback on Version 1 and detailed correspondence with Prof. Joan-Ramon Daban (Universitat Autonoma de Barcelona) on the relationship between the Metaphase Chromatin Unit (MCU) framework and the multilayer chromatin model. Framing. Scope narrowed from eukaryotic to mammalian metaphase chromosomes. The MCU is now consistently framed as an operational chromosome-scale scaling reference, not a discrete structural quantum and not a new level of the chromatin hierarchy. Figure 5 places the MCU alongside the compaction hierarchy rather than at Level 9. Reconciliation of competing structural models. New material in Introduction, Section 3.7, Section 4.1, and Conclusions integrates the Daban multilayer model with the Gibcus 2018 helical loop-array model as complementary descriptions at different structural scales, following Daban's suggestion that Hi-C-detected nested loops are compacted into the planar chromatin layers observed by microscopy. The 7.6 Mb and 250 nm MCU corresponds simultaneously to one Gibcus late-metaphase helical turn and to approximately fifteen stacked 6 nm Daban chromatin layers. The observed in vivo DNA concentration of approximately 0.17 Gb per cubic micrometre is highlighted as a physical constraint more naturally accommodated by nucleosome-nucleosome interactions between planar layers than by extended-fiber or loop-extrusion-only models. New literature integrated. Bak et al. 1979 (band splitting); Camara et al. 2024 (helical chromonema coiling across eukaryotes); Heng et al. 2013 (karyotype heterogeneity); Kramer et al. 2021 (cross-species chromosome scaling across more than 600 vertebrate and angiosperm species, discussed in a new Section 4.5); and Daban 2014, 2015, 2021, and 2024 (multilayer chromatin framework and the planar orientation of cytogenetic bands, sister chromatid exchanges, and cancer translocations). Additional caveats. Metaphase dimensions are framed as prometaphase-population averages subject to biological variability. The negative sub-500 kb quantization test is emphasized: karyometric data alone cannot distinguish a discrete structural unit from continuous proportional compaction, and finer-scale quantization (including the 0.5 Mb per 6 nm Daban layer) is not testable by cytogenetic measurement. Optical resolution limits (approximately 200 nm) are discussed in Methods. The 30 nm chromatin fiber is retained in the hierarchy figure but labeled as a historically named level rather than an in vivo confirmed architecture. No changes to underlying data, analysis pipeline, or reported quantitative results. The 33.4 nm per Mb compaction constant, the 9-of-9 microdeletion syndrome classification, the approximately 6 Mb Abbe diffraction threshold, the interactive MCU calculator, and the Zenodo archive are unchanged.

