## Supplementary material for "The Metaphase Chromatin Unit: A Novel Unit of Higher-Order Chromosome Organization in Human Mitotic Cells": Supplmentary Information

### **Contents**

Section S1. Pooled per-chromosome cytogenetic and assembly data (Table S1).

Section S2. T2T-CHM13 v2.0 arm-level reanalysis.

Section S3. Roadmap E116 chromHMM cross-validation.

Section S4. Pope 2014 Repli-Seq direct reanalysis.

Section S5. Real per-chromosome Hi-C boundary regression.

Section S6. Clinical microdeletion syndrome classification (Table S2).

Section S7. Test for integer-multiple clustering of recurrent microdeletion sizes (Figure S4).

Section S8. Cross-species predictions for mouse, rat, chimpanzee, and chicken (Figure S1).

Section S9. Complete chromatin compaction hierarchy from B-form DNA to whole chromatid (Table S3).

Section S10. Aneuploidy length predictions (Table S4).

Supplementary Figure Legends (Figures S1 to S4).

Supplementary References.

### **S1. Pooled per-chromosome cytogenetic and assembly data**

Table S1 reports the per-chromosome data used throughout the main analysis: pooled mean metaphase length from five independent cytogenetic studies of cultured peripheral lymphocytes; the inter-study standard deviation; DNA content from the GRCh38 and T2T-CHM13 v2.0 primary assemblies; cross-karyotype compaction density; and residuals from the linear fit under each assembly. Length means range from 1.576 μm (chromosome 21) to 8.208 μm (chromosome 1). The compaction density is 33.4 +/- 1.0 nm/Mb, with a coefficient of variation of 2.9 percent across all 24 chromosomes.

**Table S1.** Per-chromosome data for all 24 human chromosomes.

| **Chr** | **GRCh38 Mb** | **T2T Mb** | **Length (μm)** | **SD (μm)** | **L/Mb (nm/Mb)** | **GRCh38 resid (μm)** | **T2T resid (μm)** |
| --- | --- | --- | --- | --- | --- | --- | --- |
| 1 | 248.96 | 248.39 | 8.208 | 0.066 | 32.97 | -0.028 | +0.037 |
| 2 | 242.19 | 242.70 | 7.976 | 0.077 | 32.93 | -0.037 | -0.008 |
| 3 | 198.30 | 201.11 | 6.676 | 0.033 | 33.67 | +0.107 | +0.058 |
| 4 | 190.21 | 193.57 | 6.338 | 0.037 | 33.32 | +0.035 | -0.032 |
| 5 | 181.54 | 182.05 | 6.054 | 0.059 | 33.35 | +0.037 | +0.063 |
| 6 | 170.81 | 172.13 | 5.704 | 0.042 | 33.39 | +0.040 | +0.039 |
| 7 | 159.35 | 160.57 | 5.322 | 0.040 | 33.40 | +0.035 | +0.036 |
| 8 | 145.14 | 146.26 | 4.828 | 0.049 | 33.26 | +0.009 | +0.013 |
| 9 | 138.39 | 150.62 | 4.528 | 0.053 | 32.72 | -0.069 | -0.431 |
| 10 | 133.80 | 134.76 | 4.508 | 0.058 | 33.69 | +0.062 | +0.071 |
| 11 | 135.09 | 135.13 | 4.516 | 0.038 | 33.43 | +0.028 | +0.066 |
| 12 | 133.28 | 133.32 | 4.466 | 0.036 | 33.51 | +0.037 | +0.076 |
| 13 | 114.36 | 113.57 | 3.580 | 0.091 | 31.30 | -0.226 | -0.161 |
| 14 | 107.04 | 101.16 | 3.408 | 0.043 | 31.84 | -0.157 | +0.075 |
| 15 | 101.99 | 99.75 | 3.292 | 0.053 | 32.28 | -0.107 | +0.005 |
| 16 | 90.34 | 96.33 | 3.054 | 0.087 | 33.81 | +0.038 | -0.121 |
| 17 | 83.26 | 84.28 | 2.858 | 0.047 | 34.33 | +0.075 | +0.079 |
| 18 | 80.37 | 80.54 | 2.682 | 0.018 | 33.37 | -0.006 | +0.026 |
| 19 | 58.62 | 61.71 | 2.116 | 0.063 | 36.10 | +0.144 | +0.079 |
| 20 | 64.44 | 66.21 | 2.248 | 0.036 | 34.88 | +0.085 | +0.063 |
| 21 | 46.71 | 45.09 | 1.576 | 0.062 | 33.74 | -0.004 | +0.085 |
| 22 | 50.82 | 51.32 | 1.732 | 0.066 | 34.08 | +0.017 | +0.036 |
| X | 156.04 | 154.26 | 5.142 | 0.085 | 32.95 | -0.036 | +0.064 |
| Y | 57.23 | 62.46 | 1.846 | 0.058 | 32.26 | -0.080 | -0.216 |

### **S2. T2T-CHM13 v2.0 arm-level reanalysis**

Repeating the arm-level analysis introduced in Section 3.4 of the main text using T2T-CHM13 v2.0 centromere positions and Mb values gives a different picture from GRCh38. T2T-CHM13 correctly resolves the acrocentric short arms (10.1 to 16.7 Mb), most notably chromosome 14, whose T2T p-arm size of 11.4 Mb is 5.8 Mb smaller than the GRCh38 value of 17.2 Mb. Eight chromosomes have p-arm Mb differences greater than 1.5 Mb between assemblies. Recomputing the residual-versus-p-arm-fraction correlation with T2T values gives r = 0.24, p = 0.27, no longer statistically significant. The strong GRCh38 correlation (r = 0.65, p = 5.6e-4) was therefore partly driven by GRCh38 overestimating acrocentric p-arm Mb. The whole-chromosome T2T residuals still show the satellite over-condensation signal for chromosomes 9, 16, and Y, but a clean arm-level resolution of this signal must wait for matched arm-by-arm cytometric measurements on T2T-era data.

### **S3. Roadmap E116 chromHMM cross-validation**

We computed per-chromosome active chromatin and heterochromatin fractions from the Roadmap Epigenomics E116 GM12878 15-state chromHMM segmentation {Ernst, 2017, 29039415}. Active chromatin (states 1 to 7) correlates with the per-chromosome length residual at r = +0.48, p = 0.02 (Figure S3B); heterochromatin (states 9 and 15) correlates at r = -0.50, p = 0.014. Chromosome 19 (30.8 percent active, residual +0.144 μm) and chromosome X (87.5 percent heterochromatin in this female line, dominated by the inactive Xi) sit at opposite ends. This is independent, ChIP-seq-based confirmation of the satellite over-condensation interpretation from the GRCh38 arm-level analysis, integrating five histone-modification tracks rather than relying on a single readout.

### **S4. Pope 2014 Repli-Seq direct reanalysis**

We downloaded the segmented Repli-Seq data for GM12878 from GEO GSE53984 {Pope, 2014, 25409831} and computed the exact per-chromosome early-replication-domain (ERD) fraction. Across 23 chromosomes (Y absent in the female line), the correlation between ERD percent and length residual is r = +0.26, p = 0.24, not significant (Figure S3A). Excluding the satellite-dominated acrocentric chromosomes 13, 14, 15, 21, 22, the correlation rises to r = +0.49, p = 0.04. A multiple regression of length residual on standardized ERD percent, active chromatin percent (Section S3), and p-arm fraction yields a multiple R-squared of 0.54, with p-arm fraction contributing the largest unique R-squared (0.16), active chromatin second (0.11), and ERD third (0.06). The dominant contribution to the per-chromosome residual is satellite over-condensation captured by p-arm fraction, not replication timing per se.

### **S5. Real per-chromosome Hi-C boundary regression**

Using the actual HiCCUPS loop list (n = 9,449) and Arrowhead contact-domain list (n = 9,275) from the Rao 2014 GM12878 in situ Hi-C dataset {Rao, 2014, 25497547}, per-chromosome loop counts range from 101 (chromosome 21) to 931 (chromosome 1). Regressing metaphase length on the measured per-chromosome boundary count (excluding chromosome Y, absent from the female cell line) gives a slope of 8.78 nm per HiCCUPS loop (R-squared = 0.841, p = 7.6e-10) and 8.94 nm per Arrowhead contact domain (R-squared = 0.777, p = 2.7e-8). Mb remains the best predictor at R-squared = 0.998; nevertheless, per-chromosome boundary density (coefficient of variation 27 percent across the karyotype) correlates significantly with the length residual at r = 0.55, p = 6.6e-3. Chromosomes with higher Hi-C boundary density per Mb (chromosome 19 at 5.4 domains/Mb) have positive length residuals; lower density (chromosomes 13, 14, 15) have negative residuals. This provides a Hi-C-based test of per-chromosome fine-grained variation in compaction.

### **S6. Clinical microdeletion syndrome classification**

The approximately 6 Mb optical-resolution threshold derived in Section 3.8 of the main text, numerically close to one MCU but conceptually distinct, makes specific quantitative predictions for nine well-characterized contiguous-gene microdeletion syndromes (Table S2, Figure S2A). The framework correctly classifies all nine as karyotype-visible versus FISH-required.

**Table S2.** Predicted versus empirical karyotype detection for 9 microdeletion syndromes.

| **Syndrome (locus)** | **Size (Mb)** | **Predicted nm** | **Predicted detection** | **Empirical detection** |
| --- | --- | --- | --- | --- |
| Williams (7q11.23) | 1.6 | 53 | FISH-required | FISH-detected |
| 22q11.2 (DiGeorge) | 3.0 | 99 | FISH-required | FISH-detected |
| Smith-Magenis (17p11.2) | 3.7 | 122 | FISH-required | FISH-detected |
| 1p36 deletion | 3.5 | 115 | FISH-required | FISH-detected |
| Angelman / PWS (15q11-13) | 5.6 | 184 | Borderline | Borderline |
| 22q11.2 atypical | 6.5 | 214 | Karyotype-visible | Karyotype-visible |
| Jacobsen (11q23.3) | 11.0 | 362 | Karyotype-visible | Karyotype-visible |
| Wolf-Hirschhorn (4p16.3) | 25.0 | 823 | Karyotype-visible | Karyotype-visible |
| Cri-du-chat (5p15.2) | 30.0 | 987 | Karyotype-visible | Karyotype-visible |

### **S7. Test for integer-multiple clustering of recurrent microdeletion sizes**

We tested 26 NAHR-mediated recurrent microdeletion and microduplication syndromes (sizes 0.22 to 6.6 Mb) for clustering at integer multiples of candidate domain sizes from 100 kb to 1200 kb (Figure S4). For each candidate unit size, we computed the mean absolute residual of each CNV from its nearest integer multiple of the unit, normalized by the unit size; a quantized distribution should yield residuals near zero, whereas a uniform distribution yields a baseline of 0.25. Observed residuals lie between 0.20 and 0.34 across all tested unit sizes, statistically indistinguishable from the random baseline. No specific unit size produces quantized clustering. The strongest clustering occurs at characteristic LCR-driven sizes (1.4 to 1.7 Mb, 5 syndromes; 3.0 to 3.7 Mb, 4 syndromes), reflecting the underlying biology that NAHR-mediated CNV boundaries are positioned by segmental duplications, not by chromatin domain boundaries {Stankiewicz, 2010, 20546912; Carvalho, 2016, 27411629}. The MCU framework therefore predicts the cytogenetic detectability of CNVs of any size via the approximately 6 Mb optical-resolution threshold, without predicting their size distribution.

### **S8. Cross-species predictions for mouse, rat, chimpanzee, and chicken**

Applying the human regression to GRCm39 mouse, mRatBN7.2 rat, panTro6 chimpanzee, and galGal6 chicken assemblies yields predicted metaphase lengths that fall within published cytogenetic ranges (Figure S1). Mouse predictions span 2.06 to 6.47 μm, within the published range of 2 to 7 μm {Cowell, 1984, 6745006}. Chimpanzee predictions span 0.91 to 8.21 μm; note that the panTro6 chr 2A + 2B total of 362 Mb is inflated by approximately 50 percent relative to human chr 2 owing to known pericentromeric satellite expansion in panTro6 {Kronenberg, 2018, 29880755}, rather than reflecting real biological size difference. Chicken macrochromosomes (greater than 20 Mb each, n = 13) yield 0.5 to 6.55 μm, consistent with published macrochromosome lengths; microchromosomes (less than 20 Mb, n = 17) yield 0.14 to 0.70 μm, near the cytogenetic resolution limit. These cross-species predictions are framed as falsifiability checks rather than independent slope tests, since the human regression slope is applied directly to non-human Mb values without an independent species-specific calibration.

### **S9. Complete chromatin compaction hierarchy**

Table S3 reports the complete chromatin compaction hierarchy from the 2 nm DNA double helix to the whole metaphase chromatid in 10 hierarchical levels. The MCU defined in Section 3.7 of the main text (Level 9) closes the previously missing quantitative gap between mitotic loops (Levels 6 and 7) and whole chromosomes (Level 10). The full hierarchy is also presented graphically in main-text Figure 5.

**Table S3.** Complete chromatin compaction hierarchy from B-form DNA to whole chromatid.

| **Level** | **Structure** | **Dimensions / count** | **Cumulative compaction** |
| --- | --- | --- | --- |
| L1 | DNA double helix (B-form) | 2 nm wide; 0.34 nm/bp | 1× (baseline) |
| L2 | Nucleosome | 11 nm wide; 147 bp + 50 bp linker | ~6× |
| L3 | 10 nm nucleosomal fiber | ~200 bp / nucleosome | ~7× |
| L4 | 30 nm chromatin fiber | ~6 nucleosomes / turn; ~1.2 kb / turn | ~40× |
| L5 | Sub-loop / contact domain | 185 kb median; ~10,000 per genome | ~250× |
| L6 | Prophase loop | 60 kb median; ~50,000 per genome | ~500× |
| L7 | Prometaphase nested loops | 80 kb inner / 400 kb outer | ~2,500× |
| L8 | TAD (Dixon 2012) | 880 kb median; ~2,200 per genome | analogous |
| L9 | Metaphase Chromatin Unit (MCU) | 7.6 Mb; 0.25 μm axial; ~150 loops; 406 per haploid | ~10,000× |
| L10 | Whole metaphase chromatid | ~700 nm × 1.6 to 8.2 μm; 6 to 33 MCUs | ~10,000× |

### **S10. Aneuploidy length predictions**

The 33.4 nm/Mb compaction constant translates directly into testable cytometric predictions for common viable human aneuploidies (Table S4, Figure S2B). Trisomy 21 adds 1.58 μm of metaphase chromatin per cell (1.54 percent of total haploid mitotic chromosome length); trisomy 18 adds 2.68 μm; trisomy 13 adds 3.58 μm. The sex-chromosome aneuploidies XXX, XXY (Klinefelter), and XYY add 5.13, 5.13, and 1.85 μm respectively. Monosomy X (Turner syndrome) removes 5.13 μm. These predictions provide an upper bound on the cytometric detectability of mosaic aneuploidies: at 5 percent mosaicism the average length increase per cell falls within the inter-study SD (approximately 50 nm), suggesting that mosaic aneuploidies below 10 percent are difficult to detect by total-chromatin imaging alone.

**Table S4.** Aneuploidy length predictions from the 33 nm/Mb compaction constant.

| **Aneuploidy** | **Affected chromosome** | **MCU change** | **Length change (μm)** |
| --- | --- | --- | --- |
| Trisomy 13 (Patau) | chr 13 (+114.4 Mb) | +15.0 | +3.58 |
| Trisomy 18 (Edwards) | chr 18 (+80.4 Mb) | +10.6 | +2.68 |
| Trisomy 21 (Down) | chr 21 (+46.7 Mb) | +6.1 | +1.58 |
| Klinefelter (XXY) | +X (156.0 Mb) | +20.5 | +5.13 |
| XYY | +Y (57.2 Mb) | +7.5 | +1.85 |
| Triple X (XXX) | +X (156.0 Mb) | +20.5 | +5.13 |
| Turner (X0) | -X (156.0 Mb) | -20.5 | -5.13 |

### **Supplementary Figure Legends**

**Figure S1.** Cross-species predictions. Measured human metaphase lengths (blue circles, GRCh38) compared with predicted lengths for mouse (red squares, GRCm39), rat (purple diamonds, mRatBN7.2), chimpanzee (green triangles, panTro6), and chicken (orange crosses, galGal6) obtained by applying the human regression slope. Dashed line: human OLS fit. All non-human predictions fall within published cytogenetic ranges for those species.

**Figure S2.** (A) Microdeletion syndrome predicted length effects relative to the Abbe diffraction limit (200 nm); red = FISH-required, green = karyotype-visible. (B) Aneuploidy length predictions for common viable trisomies and sex aneuploidies. (C) Predicted chromothripsis derivative-chromosome length as a function of fragment loss percentage; material-conservative shattering remains within the Abbe-bounded undetectable zone. (D) Log-log polymer scaling: rigid rod (exponent ~1.0) versus self-avoiding walk (0.6) and fractal globule (0.33) models; data reject all globule models at more than 30 standard deviations.

**Figure S3.** (A) Per-chromosome length residual versus Pope 2014 GM12878 ERD percent (Section S4; r = 0.26, p = 0.24, not significant). (B) Per-chromosome length residual versus Roadmap E116 chromHMM active chromatin percent (Section S3; r = 0.48, p = 0.02). (C) Variance partitioning of total length variance: 99.81 percent Mb-driven, 0.09 percent inter-study measurement, 0.10 percent arm-level + chromHMM + ERD, 0.09 percent truly unexplained. (D) Independent cross-validation of the 5-study cytogenetic mean against Bender and Gooch 1962 relative-length measurements: r = 0.9995, slope = 1.02 across six decades of methodological evolution.

**Figure S4.** Test for integer-multiple clustering of recurrent microdeletion sizes (Section S7). (A) Size distribution of 33 recurrent CNV syndromes catalogued (26 of which are NAHR-mediated and form the basis of the integer-clustering test in panel C); vertical lines mark candidate chromatin domain unit sizes. (B) Cumulative distribution on log axis showing the relation to the Abbe limit, Dixon TAD, and 1 MCU. (C) Mean absolute residual from nearest integer multiple, normalized by unit size, across candidate unit sizes from 100 to 1200 kb. All values sit between 0.20 and 0.34, indistinguishable from the random baseline at 0.25. (D) Recurrent microdeletion sizes (red bars) overlaid on the chromatin hierarchy from Rao contact domain through Dixon TAD to 1 MCU.
