## Supplementary material for "The Metaphase Chromatin Unit: A Novel Unit of Higher-Order Chromosome Organization in Human Mitotic Cells": Online Calculator

Metaphase Chromatin Unit (MCU) Calculator — Goyal & Goyal 2026


### Metaphase Chromatin Unit (MCU) Calculator

Supplementary tool for Goyal & Goyal 2026.
Translate between DNA content (Mb), metaphase length (μm),
MCUs, Hi-C boundaries, and cytogenetic detection thresholds.

**Conversion constants used (this manuscript):**  
 • 1 MCU = 7.6 Mb DNA = 0.25 μm metaphase axial length  
 • Linear compaction = 33.4 nm/Mb (95% CI [32.3, 33.5])  
 • Abbe diffraction limit ≈ 196 nm at NA 1.4, λ 550 nm = 6.0 Mb threshold  
 • 1 MCU contains ≈ 41 Rao 2014 contact domains, ≈ 9 Dixon 2012 TADs, ≈ 150 Gibcus mitotic loops

#### Universal input — enter any value, others update automatically

DNA content (Mb):

Mb
= 13.16 MCU

Metaphase length (μm):

μm
= 13.16 MCU

Metaphase length (nm):

nm

MCU count:

MCU
= 1 Gibcus helical turn each

Rao contact domains (185 kb):

domains

Dixon TADs (880 kb):

TADs

Gibcus mitotic outer loops (400 kb):

loops

#### Cytogenetic CNV detectability

Predicted: above 6.0 Mb threshold — karyotypically visible

A 100 Mb DNA segment contributes ~3340 nm of metaphase length, well above the 196 nm Abbe diffraction limit.
This is a "visible" structural change on standard 400-band karyotype.

#### Reference: per-chromosome MCU breakdown (GRCh38)

| Chr | Mb | MCU | Predicted L (μm) | Observed L (μm) |
| --- | --- | --- | --- | --- |

#### Aneuploidy length predictions

| Aneuploidy | Affected chromosome | MCU change | Length change (μm) |
| --- | --- | --- | --- |
| Trisomy 13 (Patau) | chr13 (+114.4 Mb) | +15.0 | +3.76 |
| Trisomy 18 (Edwards) | chr18 (+80.4 Mb) | +10.6 | +2.64 |
| Trisomy 21 (Down) | chr21 (+46.7 Mb) | +6.1 | +1.54 |
| Klinefelter (XXY) | +X (156.0 Mb) | +20.5 | +5.13 |
| Triple X | +X (156.0 Mb) | +20.5 | +5.13 |
| XYY | +Y (57.2 Mb) | +7.5 | +1.88 |
| Turner (X0) | −X (156.0 Mb) | −20.5 | −5.13 |

#### Microdeletion syndrome quick reference

| Syndrome | Mb | MCU | nm | Detection |
| --- | --- | --- | --- | --- |
| Williams (7q11.23) | 1.6 | 0.21 | 53 | FISH |
| 22q11.2 (DiGeorge) | 3.0 | 0.39 | 99 | FISH |
| 1p36 deletion | 3.5 | 0.46 | 115 | FISH |
| Smith-Magenis (17p11.2) | 3.7 | 0.49 | 121 | FISH |
| Angelman/PWS (15q11-13) | 5.6 | 0.74 | 184 | Borderline |
| 22q11.2 atypical | 6.5 | 0.86 | 213 | Karyotype |
| Jacobsen (11q23.3) | 11 | 1.45 | 361 | Karyotype |
| Wolf-Hirschhorn (4p16.3) | 25 | 3.29 | 821 | Karyotype |
| Cri-du-chat (5p15.2) | 30 | 3.95 | 985 | Karyotype |

Goyal & Goyal 2026. This calculator is intended as a quantitative aid for clinical cytogenetics,
Hi-C interpretation, and chromosome biology. All conversions follow the manuscript's
empirical 33.4 nm/Mb scaling and the 1 MCU = 7.6 Mb = 0.25 μm definition.
