## Supplementary figures and images for "The Metaphase Chromatin Unit: A Novel Unit of Higher-Order Chromosome Organization in Human Mitotic Cells"

### Supplemental Figure 1

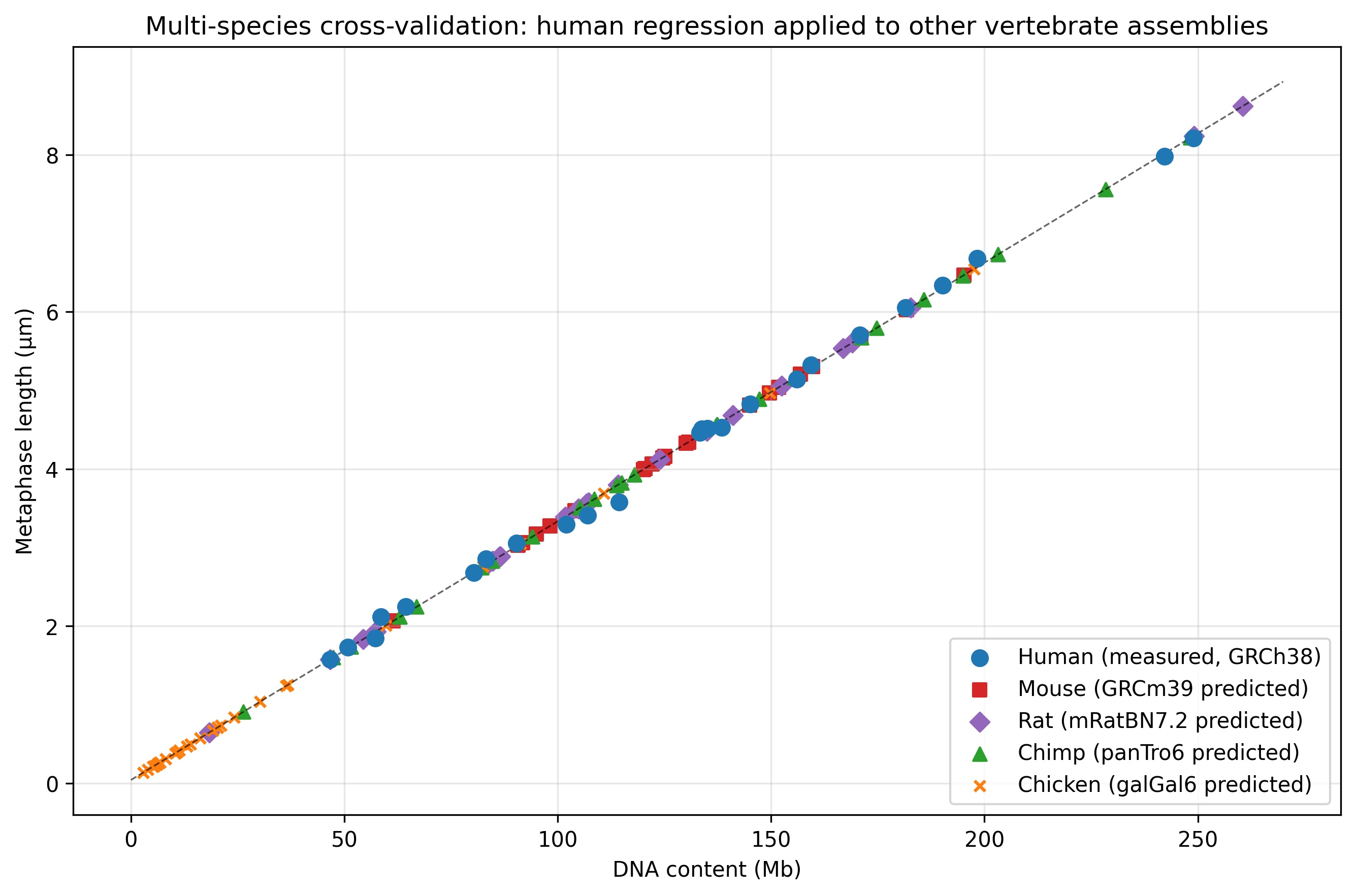

### Supplemental Figure 2

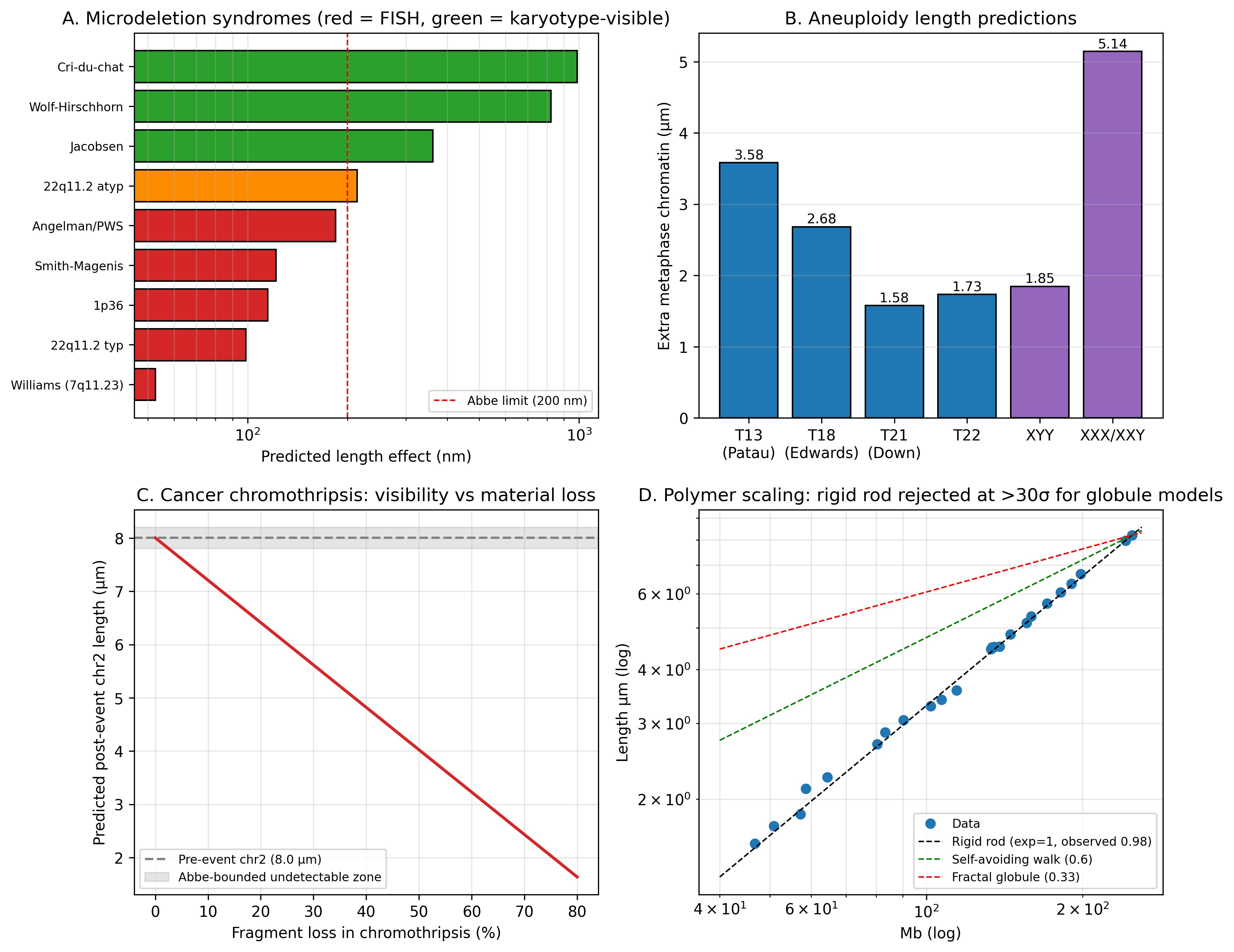

### Supplemental Figure 3

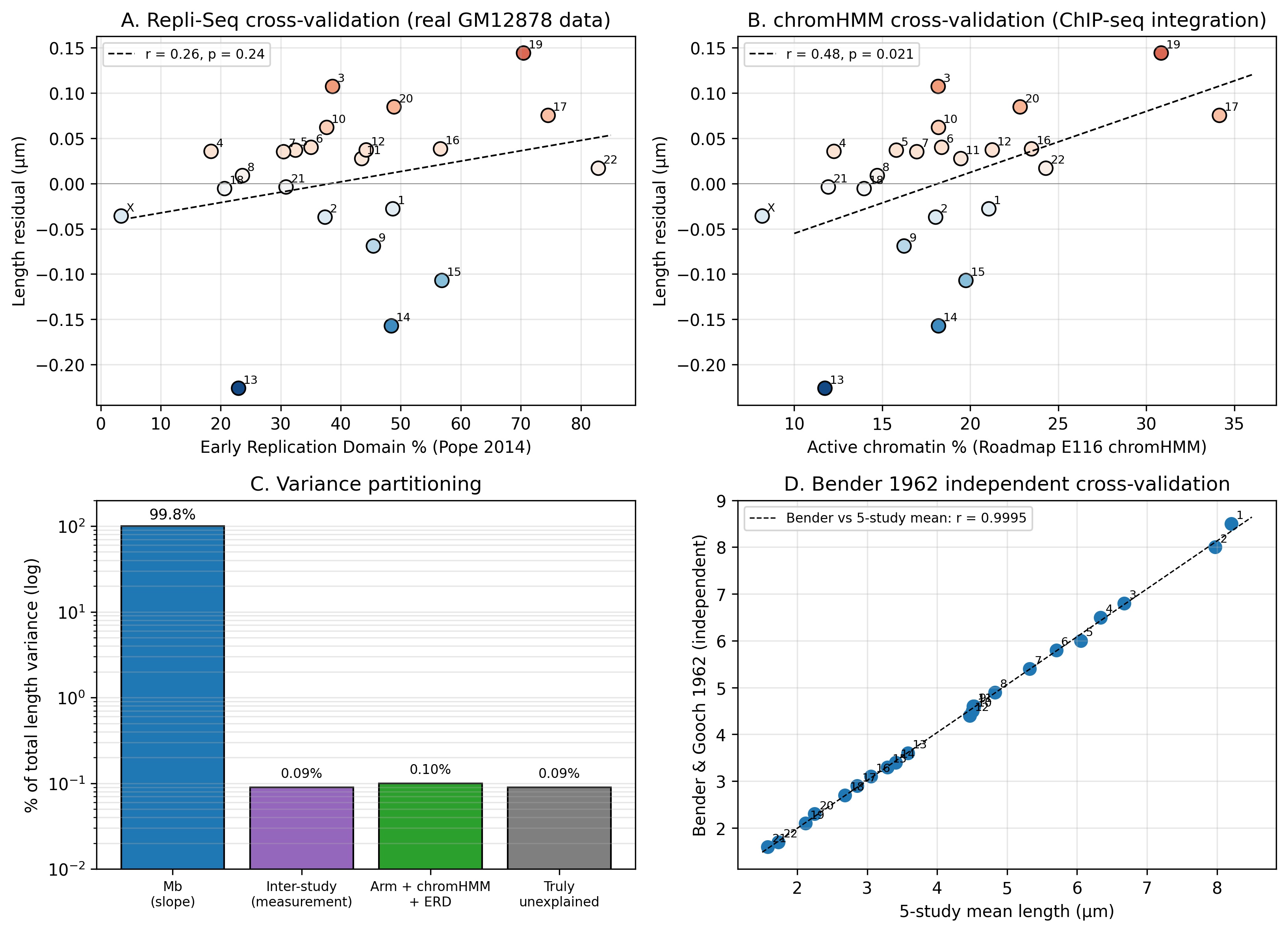

### Supplemental Figure 4

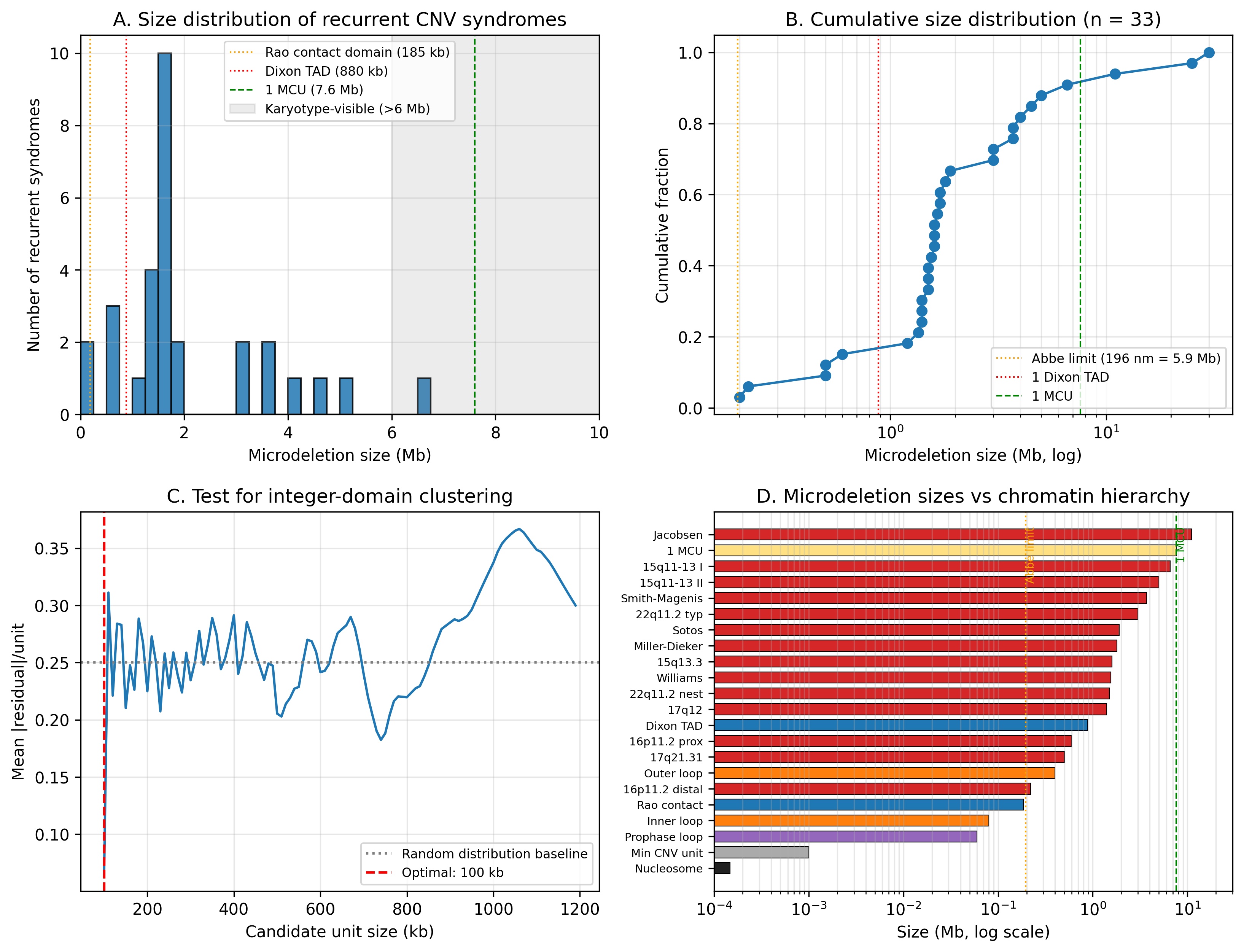
